# The effects of nano– and microplastic ingestion on the survivorship and reproduction of *Aedes aegypti* (L.) and *Aedes albopictus* (Skuse)

**DOI:** 10.1101/2023.06.23.546347

**Authors:** Gabriella McConnel, Jordann Lawson, Jaclyn E. Cañas-Carrell, Corey L. Brelsfoard

**Affiliations:** *Department of Biological Sciences, 2901 Main St., Texas Tech University, Lubbock, TX 79409, United States of America; Department of Environmental Toxicology, 1207 S. Gilbert Dr., Texas Tech University, Lubbock, TX 79416, United States of America

**Keywords:** plastic pollution, toxicology, Culicidae, larval habitat, oviposition, polystyrene

## Abstract

Microplastics (MPs) and nanoplastics (NPs) are pervasive environmental pollutants that raise concerns due to their potential impact on organisms across different trophic levels. While the effects of MPs on aquatic organisms have been extensively studied, their impacts on terrestrial organisms, mainly insects, still need to be explored. This study investigates the effects of MP and NP ingestion on the survivorship and reproduction of two medically important mosquito species, *Aedes aegypti* (L.), and *Ae. albopictus* (Skuse). Larval and pupal survivorship were not significantly affected by particle size or concentration, but there was a reduction of *Ae. aegypti* pupal survivorship associated with the ingestion of 0.03 µm NPs. Results also suggest that ingesting 0.03 µm NPs reduced egg production in both mosquito species. However, there was little impact of 0.03 NP and 1.0 µm MP ingestion on adult survivorship and longevity. To further investigate the effects of MP ingestion on mosquito fitness, we also examined the effects of lab generated MPs of varying shape, size, and plastic polymer type on *Ae. aegypti* immature and adult survivorship. The data suggests that the polymer type and shape did not impact *Ae. aegypti* immature or adult survivorship. These findings highlight the potential consequences and the need to investigate further the ecological and potential public health implications of MP and NP ingestion by mosquitoes.

## 1. Introduction

Micro-(MPs) and nanoplastics (NPs) are ubiquitous in the environment and have become a growing concern due to their potential impacts on organisms at various trophic levels. MPs are defined as plastic particles < 5 mm, and NPs as < 1 µm in size. MPs and NPs are often produced via the production of plastic pellets for plastic product manufacturing and are found in personal care and other cosmetic products (Cortes-Arriagada et al., 2023; Duis and Coors, 2016; Pinheiro et al., 2023; Shahsavaripour et al., 2023). MPs and NPs can also be formed via the degradation of plasticized products via UV light exposure, abrasion, and weathering (Fadare et al., 2020; Huang et al., 2022; Liu et al., 2023; Zhang et al., 2021). Once micro- and nanoplastics (MNPs) are formed or enter the environment they are difficult to remove from substrates and are often ingested by vertebrates and invertebrates in a variety of aquatic and terrestrial ecosystems (Carrillo et al., 2023; Clark et al., 2016; Cohen-Sanchez et al., 2023; Cole et al., 2019; Cole et al., 2020; Elizalde-Velazquez et al., 2020; Liu et al., 2022; Malafaia et al., 2020; Rochman et al., 2017; Silva et al., 2021).

Previous work on MPs has focused on examining the underlying negative physiological effects on aquatic organisms and until recently the effects of MPs and NPs on terrestrial organisms has been relatively underexplored (Elizalde-Velazquez et al., 2020; Lu. et al., 2018). Particularly, there is a knowledge gap of the effects of MP ingestion regarding terrestrial invertebrates such as insects. Recent work has demonstrated that the ingestion of irregularly shaped MPs by the midge *Chironomus riparius* (Meigen) resulted in the accumulation of MPs in larval tissues, which ultimately resulted in a reduction in growth and emergence rates (Silva et al., 2019). MNPs can vary greatly in size and composition. MP size dependent gut damage has previously been demonstrated in *D. melanogaster* (Meigen) (Zhang et al., 2020). The composition of MNPs may also be an important factor that influences the effects of MNP ingestion in invertebrates. Plastics are complex polymers that are typically combined with additives that improve functionality by giving plastics durability, flexibility, and other properties. These chemical additives in different polymer types are often weakly bound to the plastic polymers and can leach into the surrounding substrate. For example, cadmium and other related compounds are commonly used as pigments and stabilizers in polyvinyl chloride, acrylonitrile butadiene styrene, and polystyrene (Liu et al., 2020; Turner, 2019). Additionally, the tendency of these plastics to adsorb persistent organic pollutants (POPs) and release them under certain conditions adds another layer to the problem. Investigation on how these plastics may increase or decrease the delivery of these POPs to organisms post ingestion is still underway (Hahladakis et al., 2018). Results from these studies suggest there may be a varying effect associated with the size and shape of the MP or NP as well as the chemical composition of the polymer, suggesting the need to investigate the effects of MP size, shape, and composition on life history traits in insects.

Examining the effects of MNPs on mosquitoes is of particular interest as they are important vectors of multiple viruses that cause human disease, such as dengue fever, chikungunya, and Zika virus (Lin et al., 2023; Mercier et al., 2022; Novelo et al., 2023; Severini et al., 2018; Zimler and Alto, 2023). Additionally, mosquitoes provide an appropriate system to study the effects of MNPs in part due to the indiscriminate filter-feeding behavior of larva and that they are likely to feed on MPs in aquatic habitats (Merritt et al., 1992). Furthermore, container inhabiting mosquitoes are likely to be exposed to MNPs due to many preferred oviposition and larval habitats consisting of artificial man-made containers that are often composed of plastic (Champion and Vitek, 2014).

Here we investigated the effects of MP and NP ingestion on the survivorship and reproduction of two medically important mosquito species, *Aedes aegypti* (L.) and *Aedes albopictus* (Skuse). Specifically, we quantified the effects of MP and NP ingestion on mosquito survival and reproductive output. Furthermore, we also investigated the effect of irregularly shaped MPs of different plastic polymer types on survivorship and reproduction of *Ae. aegypti*. The results are discussed in relation to gaining an understanding of the impacts of MNPs on mosquito populations and on pathogen transmission.

## 2. Materials and Methods

### 2.1 Mosquito rearing

*Aedes aegypti* (Rockefeller strain) and *Aedes albopictus* colonies generated from eggs collected in Lubbock, TX, USA were used for the experiments. Mosquitoes were reared in incubators maintained at 28°C ± 1°C and 80 ± 5% RH and a 16:8 light:dark cycle. Eggs were collected in 140 mL cups lined with #76 heavy weight seed germination paper (Anchor Paper Co., MN, USA), and stored in an incubator for approximately two weeks prior to hatching. Eggs were hatched in a 1:1 ratio of a hatching solution: DI water. The hatching solution consisted of 0.03 g/L of liver powder solution inoculated with a 1 mL water sample collected from *Aedes* larval habitat in Lubbock, TX, USA. Larvae were fed a 60 g/L bovine liver powder (MP Biomedicals, Santa Ana, CA) slurry ad libitum.

### 2.2 Effects of NP and MP ingestion on Ae. aegypti and Ae. albopictus life history traits

Using a 3 mL glass transfer pipet 100 1^st^ instar larvae were placed in a 20.3 x 20.3 x 7 cm, Pyrex glass baking tray (Corning Inc., Corning NY) containing 300 mL 1:1 solution of hatching solution: DI water and varying concentrations of 1.0 µm carboxylate-modified polystyrene beads (0, 50, 100, and 10,000 beads /mL; density 1.050 g/cm^3^, excitation 470 nm, emission 505 nm) (Sigma-Aldrich, Munich, Germany). The aforementioned experiments were also repeated using 0.03 µm fluorescent green carboxylate-modified polystyrene beads (density 1.050 g/cm^3^, excitation 470 nm, emission 505 nm) (Sigma-Aldrich, Munich, Germany). MP bead types were characterized using FEI Quanta 600 field-emission scanning electron microscope (SEM) and X-ray diffraction (XRD) using previously described methods (Edwards et al., 2023) (Supplementary Figure 1). Concentrations of stock solutions of NP and MP beads were calculated using a hemocytometer and a Leica DM IL microscope with a green fluorescent protein (GFP) filter (Leica microsystems, Wetzlar, Germany) with an attached CoolLED pE-300 ultraviolet light system (Andover, SP10 5NY, United Kingdom).

Rearing pans were monitored daily after the first pupal emergence. Pupae were counted and transferred to 150 mL glass beakers containing 50 mL of DI water, which were placed in 60 cm^3^ cages. All adults were provided with 10% sucrose solution ad libitum. Four days after adult emergence, mosquitoes were provided a blood meal three times per week for three weeks using an artificial feeding system placed on each cage. Once a week, eggs were collected, counted, and oviposition cups replaced. Cages were monitored for adult mortality every 24-48 hrs for 40 days from emergence of the first adult mosquito. Each treatment type consisted of three replicates and untreated controls were replicated six times. Larval survivorship was calculated as the proportion of larvae that fully developed to pupae. Pupal survivorship was calculated as the proportion of pupae that emerged into adult mosquitoes.

### 2.3 Effect of ingestion of irregularly shaped MPs on survivorship of Ae. aegypti

Irregularly shaped MPs were lab generated by grating polyethylene terephthalate (PET), high-density polyethylene (HDPE), polypropylene (PP) and nylon fibers (NF) by grating the plastic types on a smooth cut metal file. Each MP type was imaged using a Leica S9 stereomicroscope (Leica microsystems, Wetzlar, Germany) with an attached camera. MPs were then characterized by measuring the diameter at the widest point for 10 different MP particles of each plastic type. MP measurements were conducted using ImageJ (Schneider et al., 2012).

*Ae. Aegypti* eggs were allowed to hatch for 45 min and the 50 first instar larvae were transferred to 500 mL glass beakers containing 200 mL of DI water and 0.5 mg of each MP type. A control with no MPs was also included in the experimental design. Each beaker was covered with aluminum foil and placed in an incubator. Larvae were fed a 60g/L liver powder:DI water slurry ad libitum. Pupae were transferred to a 50 mL glass beaker containing ∼15 mL of DI water and placed in an appropriately labeled cage. Adults were provided a 10% sucrose solution ad libitum. Cages were monitored for approximately three times a week for nine weeks for the presence of dead adults. The sex of each dead adult was recorded to determine adult survivorship rates. Three replicates were completed for each MP type and untreated controls.

### 2.4 Statistical Methods

A one-way ANOVA was used to determine differences between larval and pupal survivorship and adult emergence rates after ingestion of different MP types, sizes, and concentrations. All proportional values were subjected to an Arcsine square root transformation before analysis. Differences between treatments were determined using Bonferroni corrected Tukey-Kramer HSD post hoc tests. Differences in survivorship between MP treatments were examined using Kaplan-Meier survivorship curves and Log-Rank tests. All statistical analyses were performed using JMP Pro vs.16.0.0 (SAS Institute Inc., Cary, NC).

## 3. Results

### 3.1 MP effects on immature survivorship and adult emergence rates

MP size and concentration did not affect the larval survivorship of *Ae. aegypti* or *Ae. albopictus* (*Ae. aegypti*, MP size; ANOVA, F_2,21_ = 0.46, P = 0.64; MP concentration ANOVA, F_3,20_ = 1.06, P = 0.39; and *Ae. albopictus*, MP size, ANOVA, F_2,21_ = 0.18, P = 0.84; MP concentration ANOVA, F_3,20_ = 0.30, P = 0.82) (Table 1). MP concentration also did not affect *Ae. aegypti* or *Ae. albopictus* pupal survivorship (*Ae. aegypti*, ANOVA, F_3,20_ = 0.55, P = 0.65; and *Ae. albopictus,* ANOVA, F_3,20_ = 2.22, P = 0.12). However, plastic size did have a significant effect on *Ae. aegypti* pupal survivorship (ANOVA, F_2,21_ = 7.3, P = 0.004) with an observed reduction in pupal survivorship associated with the ingestion of 0.03 µm NPs, but no difference was observed in *Ae. albopictus* (ANOVA, F_2,21_ = 0.41, P = 0.67) (Table 1). There was no observed effect of plastic size or concentration on the adult emergence of *Ae. aegypti* and *Ae. albopictus* that ingested 1.0 µm MPs or 0.03 µm NPs (*Ae. aegypti*, MP size; ANOVA, F_2,21_ = 2.31, P = 0.12; MP concentration ANOVA, F_3,20_ = 0.60, P = 0.62; and *Ae. albopictus*, MP size, ANOVA, F_2,21_ = 0.06, P = 0.94; MP concentration ANOVA, F_3,20_ = 0.04, P = 0.98) (Table 1).

**Table 1.** *Ae. albopictus* and *Ae. aegypti* immature and adult survivorship after ingestion of 0.03 and 1.0 µm polystyrene MPs at different concentrations.

| Species | MP concentration (MP/mL) | MP size | N | % larval survivorship | % pupal survivorship | % adult emergence |
| --- | --- | --- | --- | --- | --- | --- |
| *Ae. albopictus* | 0 | - | 6 | 79.7 ± 3.9 | 95.3 ± 1.1 | 76.2 ± 4.0 |
| | 50 | 0.03 | 3 | 81.7 ± 3.4 | 97.4 ± 0.1 | 79.7 ± 3.4 |
| | 50 | 1.0 | 3 | 75.3 ± 3.2 | 99.1 ± 0.4 | 74.7 ± 2.9 |
| | 100 | 0.03 | 3 | 78.3 ± 3.4 | 97.8 ± 1.0 | 76.7 ± 3.7 |
| | 100 | 1.0 | 3 | 86.0 ± 3.5 | 90.2 ± 3.3 | 78.7 ± 5.5 |
| | 10,000 | 0.03 | 3 | 78.7 ± 4.6 | 95.6 ± 1.0 | 75.3 ± 3.8 |
| | 10,000 | 1.0 | 3 | 82.7 ± 2.4 | 96.2 ± 1.5 | 79.7 ± 2.9 |
| *Ae. aegypti* | 0 | - | 6 | 85.5 ± 3.5 | 89.3 ± 2.5<sup>a</sup> | 80.7 ± 3.7 |
| | 50 | 0.03 | 3 | 81.7 ± 0.3 | 74.7 ± 4.3<sup>b</sup> | 67.3 ± 1.9 |
| | 50 | 1.0 | 3 | 86.3 ± 5.2 | 96.8 ± 0.3<sup>a</sup> | 83.7 ± 5.0 |
| | 100 | 0.03 | 3 | 81.0 ± 2.3 | 87.6 ± 2.1<sup>b</sup> | 72.3 ± 3.5 |
| | 100 | 1.0 | 3 | 86.3 ± 5.5 | 96.9 ± 1.9<sup>a</sup> | 83.7 ± 4.1 |
| | 10,000 | 0.03 | 3 | 91.0 ± 3.2 | 89.6 ± 3.6<sup>a</sup> | 83.0 ± 5.5 |
| | 10,000 | 1.0 | 3 | 89.3 ± 6.3 | 91.7 ± 2.2<sup>a</sup> | 82.3 ± 4.3 |

Different superscripted letters refer to significant differences between groups in each column for each species (Tukey-Kramer post-hoc tests, $P \leq 0.05$ ).

### 3.2 NP and MP effects on female fecundity

When examining egg production per female over three gonotrophic cycles, the size of the plastic ingested, but not the concentration, had a significant impact on *Ae. albopictus* and *Ae. aegypti* fecundity (Figure 2; ANOVA, *Ae. albopictus*, bead size F_4,67_ = 5.4, P = 0.02; MP concentration F_4,67_ = 0.32, P = 0.72; *Ae. aegypti*, bead size F_4,79_ = 6.6, P = 0.01; MP concentration F_4,79_ = 0.87, P = 0.87). When *Ae. albopictus* and *Ae. aegypti* females were reared in water containing 0.03 µm polystyrene NPs they produced fewer eggs than females from control and 1.0 µm MP treatments.

**Figure 1.**
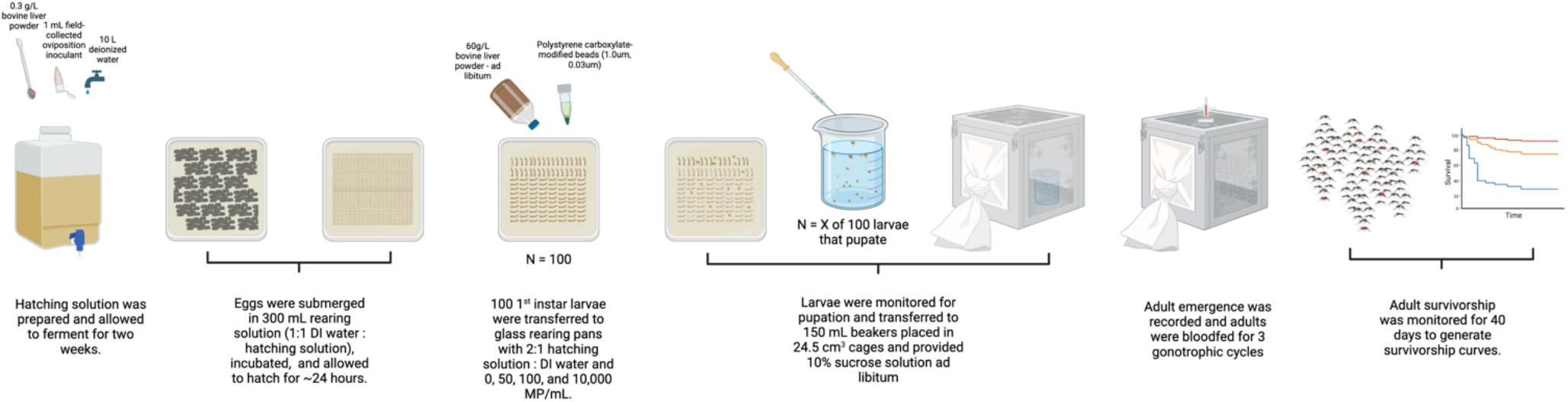
Experimental setup to measure the impact of 0.03 and 1.0 µm polystyrene MPs on the life history traits of *Ae. aegypti* and *Ae. albopictus*.

**Figure 2.**
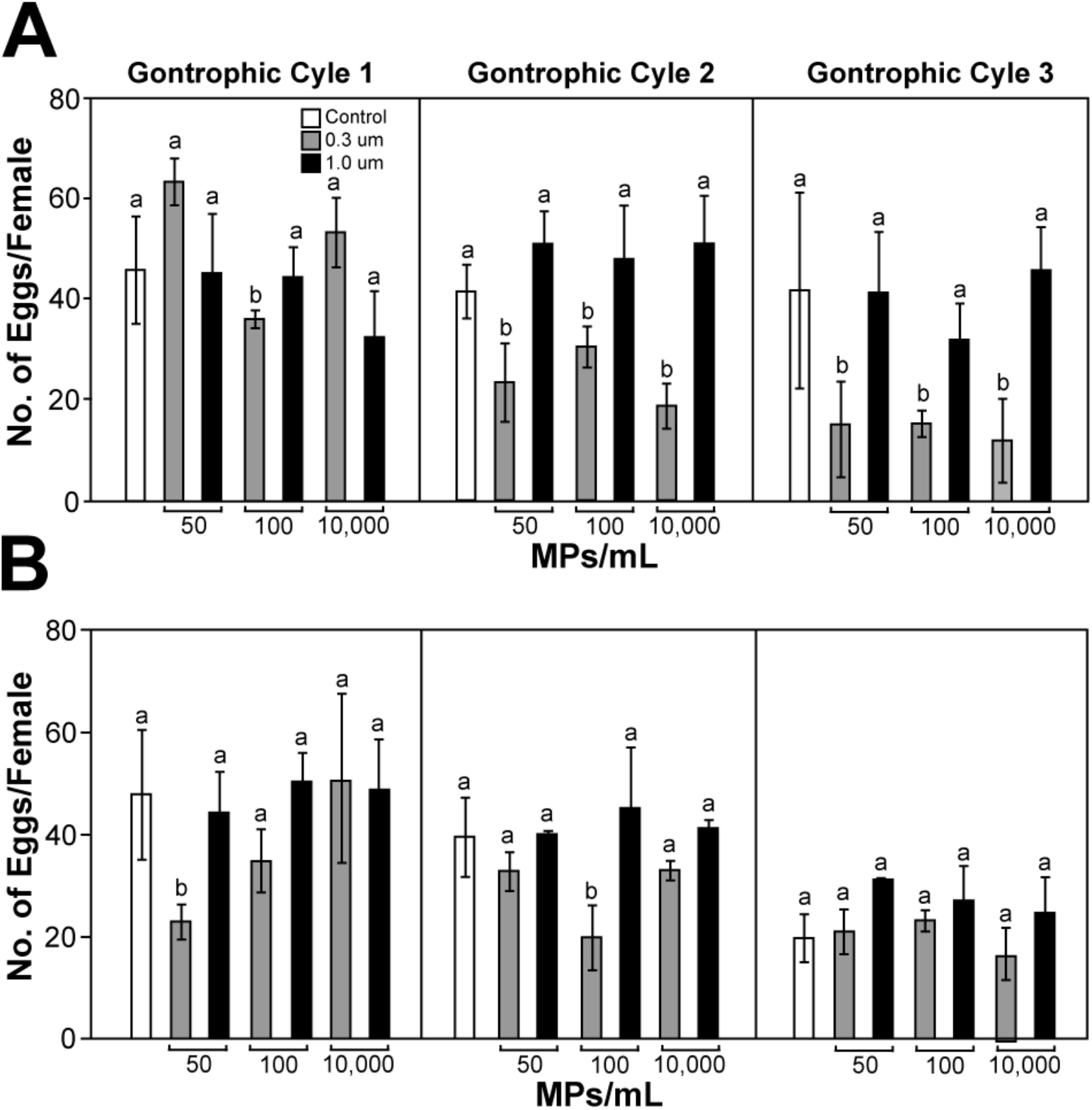
(A) *Ae. albopictus* and (B) *Ae. aegypti* fecundity over three gonotrophic cycles when exposed to 0.03 and 1.0 µm polystyrene MPs at 0, 50, 100, and 10,00 MPs/mL in rearing water. All data is displayed as the mean ± SE. Different letters above each bar coincide with statistical differences according to Bonferroni corrected post hoc tests (P < 0.01) performed for each gonotrophic cycle.

### 3.3 MP effects on adult survivorship

Kaplan-Meier survival curves were made for comparisons of *Ae. aegypti* and *Ae. albopictus* survival after ingesting different concentrations of 1.0 µm and 0.03 µm polystyrene beads. No difference between survival curves was observed in *Ae. aegypti* females that ingested 1.0 µm or 0.03 µm beads (Figure 3). However, there was an observed difference in survivorship for males that ingested 1.0 µm MPs (1.0 µm; males, Log-Rank, Chi-square = 14.1, P = 0.003; females, Log-Rank, Chi-square = 7.07, P = 0.07, 0.03 µm; males Log-Rank, Chi-square = 0.75, P = 0.86; females, Log-Rank, Chi-square = 2.74, P = 0.43). Pairwise tests with a Bonferroni correction suggested a significant difference when comparing the control group to the 1.0 µm 50 MP/mL group of *Ae. aegypti* males (Log-Rank, Chi-square = 6.8, P ≤ 0.01). No other significant differences were observed in pairwise comparisons of the MP and NP treatment groups. No difference between survival curves was observed in *Ae. albopictus* males or females ingesting 1 µm MPs or 0.03 µm NPs (1.0 µm; males, Log-Rank, Chi-square = 2.91, P = 0.40; females, Log-Rank, Chi-square = 1.48, P = 0.69, 0.03 µm; males Log-Rank, Chi-square = 1.02, P = 0.80; females, Log-Rank, Chi-square = 6.06, P = 0.11) (Figure 3). Pairwise tests with a Bonferroni correction (Bonferroni corrected p-value = 0.008) suggested there were no differences between any of the treatment groups for either bead size (P ≥ 0.008).

**Figure 3.**
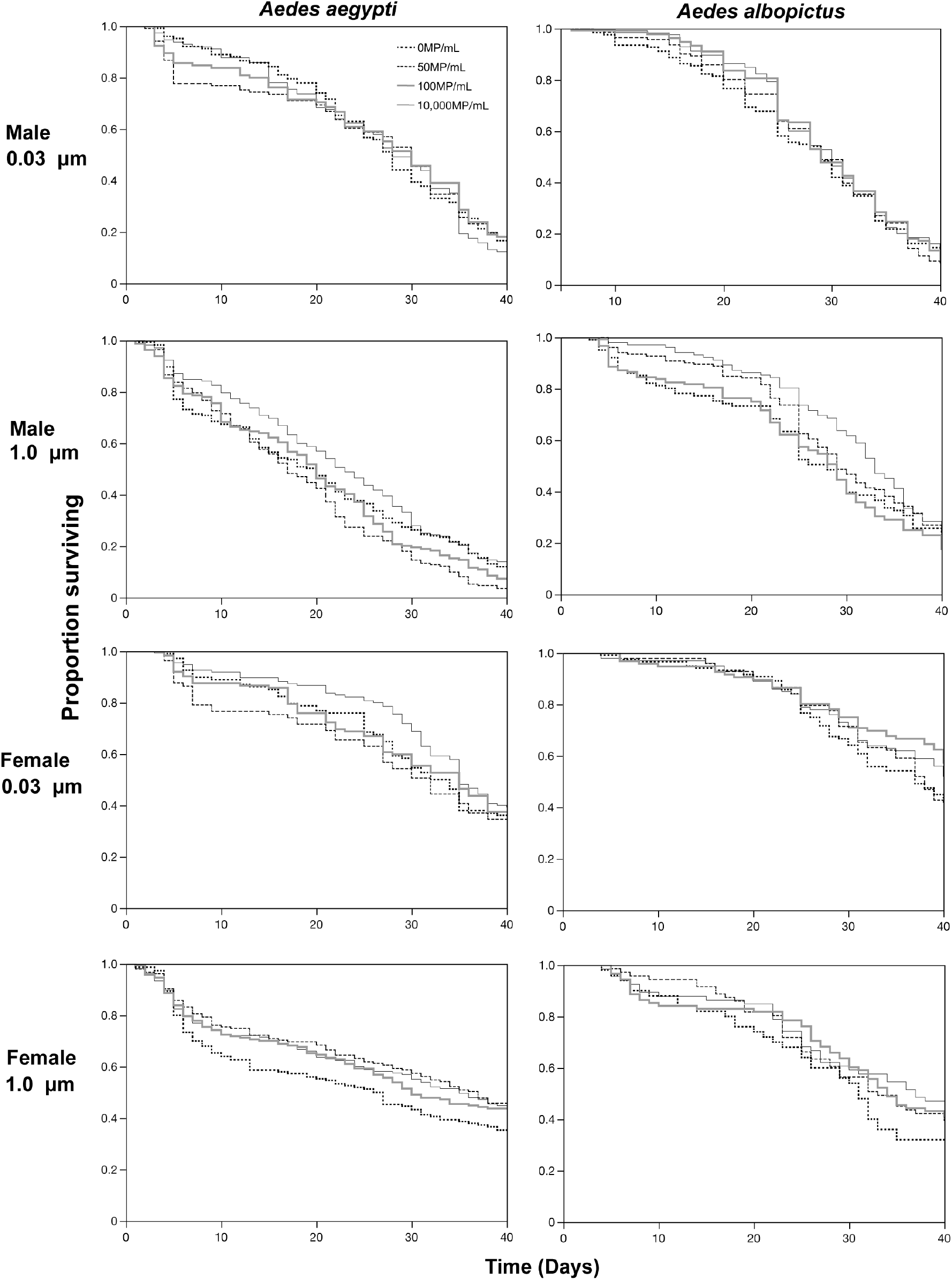
The effect of MP size and concentration on male and female *Ae. albopictus* and *Ae. aegypti* survivorship and longevity. Kaplan-Meier survivorship curves indicate little impact of the ingestion of 0.03 and 1 µm polystyrene MPs on the survivorship of *Ae. aegypti* and *Ae. albopictus*.

### 3.4 Effects of composition of irregularly shaped MPs on Aedes aegypti immature and adult survivorship

PET, PP, HDPE, and NF laboratory manufactured MPs were characterized by measuring their diameter. Mean diameter measurements are shown in Figure 4A. Larval survivorship was not affected by the ingestion of PET, PP, HDPE, and NF MPs (ANOVA, F_4,10_ = 0.30, P = 0.87) (Figure 4B). Similar to larval survivorship there was also no difference in adult emergence rates when MP treatment groups were compared to untreated controls (ANOVA, F_4,10_ = 0.18, P = 0.95) (Figure 4C). No difference in adult survivorship rates was observed when comparing MP treatments with untreated controls (Females, Log-Rank, Chi-square = 3.83, P = 0.43; Males, Log-Rank, Chi-Square = 5.71, P = 0.22) (Figure 4D). A pairwise test with a Bonferroni correction (Bonferroni corrected p-value = 0.005) showed there was a significant difference when comparing the survivorship of males that ingested PET MPs compared to the untreated control group (Log Rank, Chi-square = 5.6, P ≤ 0.001). All other pairwise comparisons of survivorship of males and females that ingested PET, PP, HDPE, and NF MPs were not significantly different (P ≥ 0.001).

**Figure 4.**
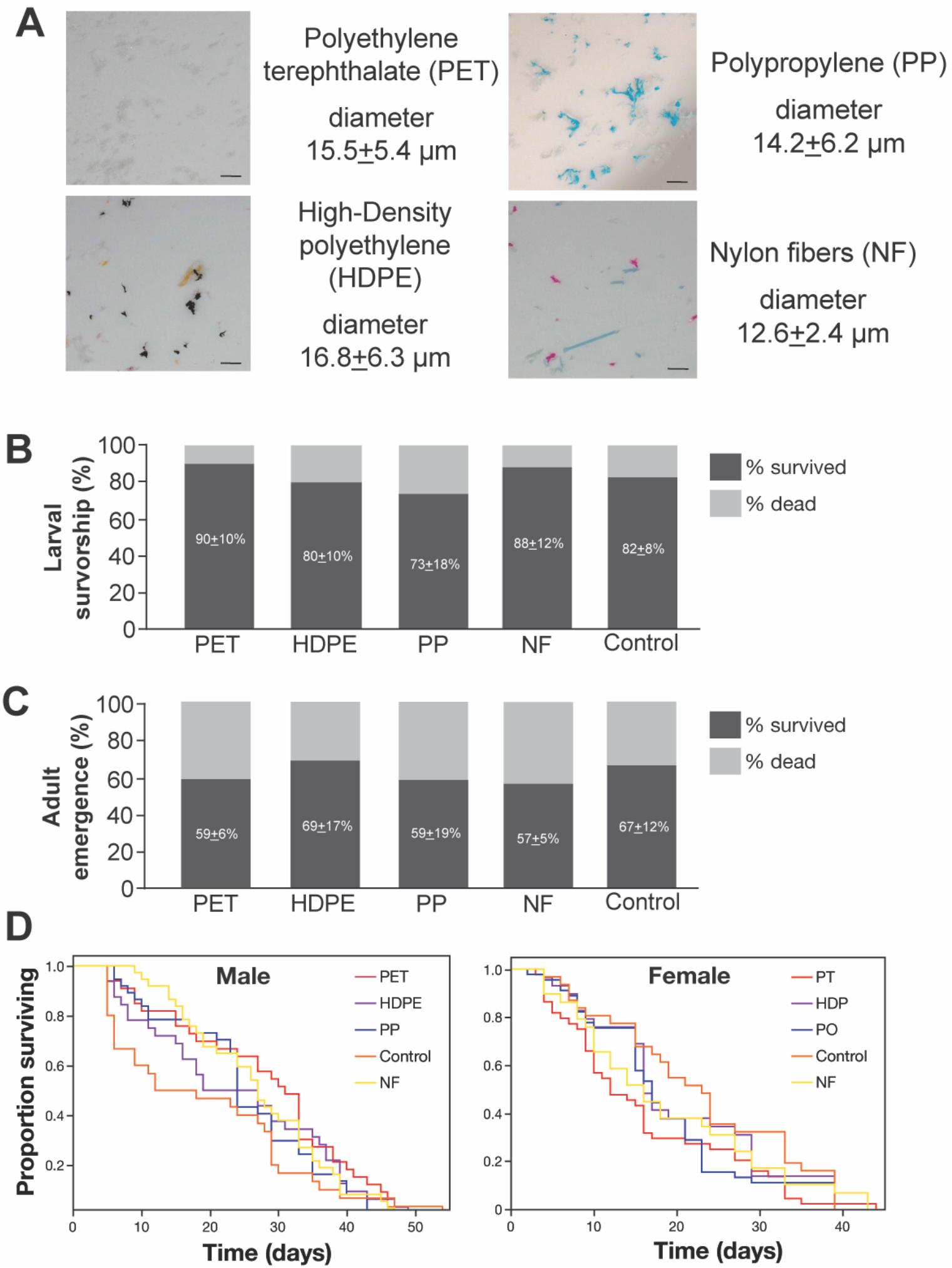
(A) images of the laboratory manufactured PET, PP, HDPE, and NF microplastic particles and associated mean ± diameter measurements, (B) The effects of MP type on larval survivorship, and (C) percent adult emergence. The numbers inside of each bar represent the mean percentage of larvae that survived ± standard error. No significant differences were observed between the different MP types for larval survivorship or adult emergence (Tukey-Kramer post hoc pairwise comparisons, P ≥ 0.05), and (D) survival curves of male and female *Ae. aegypti* reared in water containing each microplastic type. The scale bars represent 40 µm.

## 4. Discussion

Previous work demonstrated that *Ae. aegypti* and *Ae. albopictus* ingest 1.0 µm polystyrene MPs and these MPs accumulate in *Ae. aegypti* and *Ae. albopictus* guts and perturb the gut microbiome (Edwards et al., 2023). To further investigate the effects of MP and NP ingestion on *Ae. aegypti* and *Ae. albopictus*, we exposed larvae of both species to two sizes (0.03 and 1.0 µm) of polystyrene beads at three different concentrations (50, 100, and 10,000 MP/mL). The first two concentrations were considered realistic exposure concentrations that mosquitoes may experience in natural larval habitats, and the 3^rd^ concentration was included as an extreme for exposure scenarios in natural larval habitat. Our results show that the bead size and the concentration of MPs and NPs in rearing containers had little effect on immature survivorship. Only a minor effect was observed where the ingestion of 0.03 µm NPs was associated with a reduction in pupal emergence rates in *Ae. aegypti*.

The data also suggests there was little impact on adult emergence rates and adult survivorship and longevity associated with the ingestion of 0.03 NPs and 1.0 µm MPs. Significant differences in the survivorship curves were observed when comparing the 50 MP/mL treatment to untreated controls of *Ae. aegypti* males. However, this is unexpected as it was anticipated that the higher concentrations of NPs and MPs would impact adult survivorship. These observed results are consistent with previous studies where *Cx. tarsalis* and *Cx. pipiens* ingested 200 or 200,000 polystyrene MPs per mL and there was little observed effect on body size, development, and growth rate (Thormeyer and Tseng, 2023). Previous work also suggested that adults egest a large proportion of the MPs ingested as larvae shortly after emergence as adults particularly after seeking a sucrose meal, suggesting MPs may not be embedding in host gut tissues and resulting in any significant long term impacts on adult life history traits (Edwards et al., 2023). However, in this study there was an observed NP induced effect on female fecundity. Particularly, females that ingested 0.03 µm NPs had reduced fecundity compared to the 1.0 µm MP treatment group and the untreated controls. This observed effect could be due to the presence of styrene, which is used to make the polymer polystyrene. Styrene has been reported to be highly toxic to multiple cell types including epithelial cells (Hwang et al., 2020). Bisphenol A (BPA) and phthalates are commonly used plasticizers and are considered endocrine disruptors. BPA has been previously demonstrated to impact larval development time where *Culex quinquefasciatus* (Say) larval development was shortened up to 25% after exposure (Valsala and Asirvadam, 2022). Primary MNPs generated for lab use like the polystyrene beads in this study are unlikely to contain BPA, but the presence of plasticizers is something to consider for future studies that aim to bridge the gap between lab-scale exposure experiments utilizing pristine MNPs and the behavior of these MNPs in the natural environment. What is more, as the 0.03 µm beads are nanoscale, slightly smaller than virions from the genus *Flavivirus* in comparison (Simmonds et al., 2017). Due to size they may have the ability to embed themselves in insect gut tissue as well as enter the insect hemolymph and other host somatic tissues. Magni et al 2018 found even polystyrene beads at the micro-scale to concentrate in gut lumen of the freshwater zebra mussel, *Dreissena polymorpha*, before moving to hemolymph (Magni et al., 2018).This is the first study to investigate the effects nanoplastics have on mosquito physiology and this hypothesis would require additional histology studies to examine for the presence of MPs in mosquito somatic tissues.

The observed minimal impacts to host fitness particularly on survivorship and host longevity could support the hypothesis that mosquitoes act as environmental transporters for MNPs, potentially transferring MNPs to aquatic sites when visiting for oviposition or seeking mates and when seeking nectar sources. A similar hypothesis was also previously proposed and suggested that predation of mosquitoes that had previously ingested MPs could also be a route of ontogenic transfer to other invertebrate and vertebrate animals (Al-Jaibachi et al., 2019; Cui et al., 2022). Since male and female mosquitoes often visit nectar sources for a sugar meal, they could potentially translocate MNPs to nectar sources when expelling frass containing MPs after a sugar meal, providing an additional exposure source to other insects that may be visiting the same nectar source.

While there was little observed difference examining the effects of as single polymer type of uniform size, symmetry, and of only one composition type we wanted to examine the effect of irregularly shaped MPs of different polymer compositions. A similar lack of fitness effects was also observed when larvae exposed to lab generated PET, PP, HDPE, and NF microplastic particles. In this study, polymer type and shape did not appear to have any significant impacts to *Ae. aegypti* physiology. This result is in contrast to earlier work investigating the effects of irregular shaped polyethylene MPs of 17-53 µm in diameter on *C. quinquefasciatus* (Malafaia et al., 2020). In this previous study, the ingestion of irregular shaped polyethylene MPs by fourth instar larvae were associated with biochemical changes leading to impacts on metabolism, oxidative stress, and neurotoxicity. However, a potential reason for this difference in results could be due to differences in polymer type, size and morphological differences between the species as well as the developmental age of the larvae during microplastic exposure (Lima et al., 2003). In this study, to simulate more likely environmental exposure conditions, larvae were exposed as first instars, while in previous studies fourth instar larvae were exposed to MPs (Malafaia et al., 2020).

A majority of studies to date that have investigated the impacts of MPs on insects have systematically focused on examining the effects of one microplastic size and type. In nature, this maybe a completely different situation where mosquitoes are ingesting a combination of different sizes and types of plastics of different ages, where older weathered MNPs could harbor microbiota biofilms, leach additives, persistent organic pollutants like pesticides, and perhaps other xenobiotics. These results presented here offer several perspectives to continue to investigate. Specifically, it would be important to determine the fate of MNPs in mosquitoes. These studies should investigate whether NPs or MPs imbed themselves in host gut and other host somatic tissues. Furthermore, it would also be interesting to perform surveys and analyze the types, numbers, and composition of MNPs found in mosquito larval habitats, particularly artificial man-made containers as well as wild caught mosquitoes. These containers offer an unexplored portion of the plastisphere, niche microenvironments that could act as sinks or sources of these particles, and potentially behave differently from the natural freshwater and saltwater habitats where most MP descriptive studies have been performed. Ultimately, from a public health perspective it would be important to determine if the ingestion and accumulation of MPs and NPs in the mosquito adult will have any impacts on pathogen transmission.

## Acknowledgment

We would like to thank Dr. Micah J. Green and Dr. Kailash Dhondiram Arole for their help with the characterization of the polystyrene MPs used in this study. This work was funded by the US National Science Foundation grant # 2136670.

## Data availability statement

Data is available upon a reasonable request. Please contact the corresponding author Corey L. Brelsfoard with data requests.

## Supporting information

Supplemental Figure 1

## Notes

### Competing Interest Statement

The authors have declared no competing interest.

