## Supplementary figures and images for "The effects of nano– and microplastic ingestion on the survivorship and reproduction of *Aedes aegypti* (L.) and *Aedes albopictus* (Skuse)"

### Supplemental Figure 1

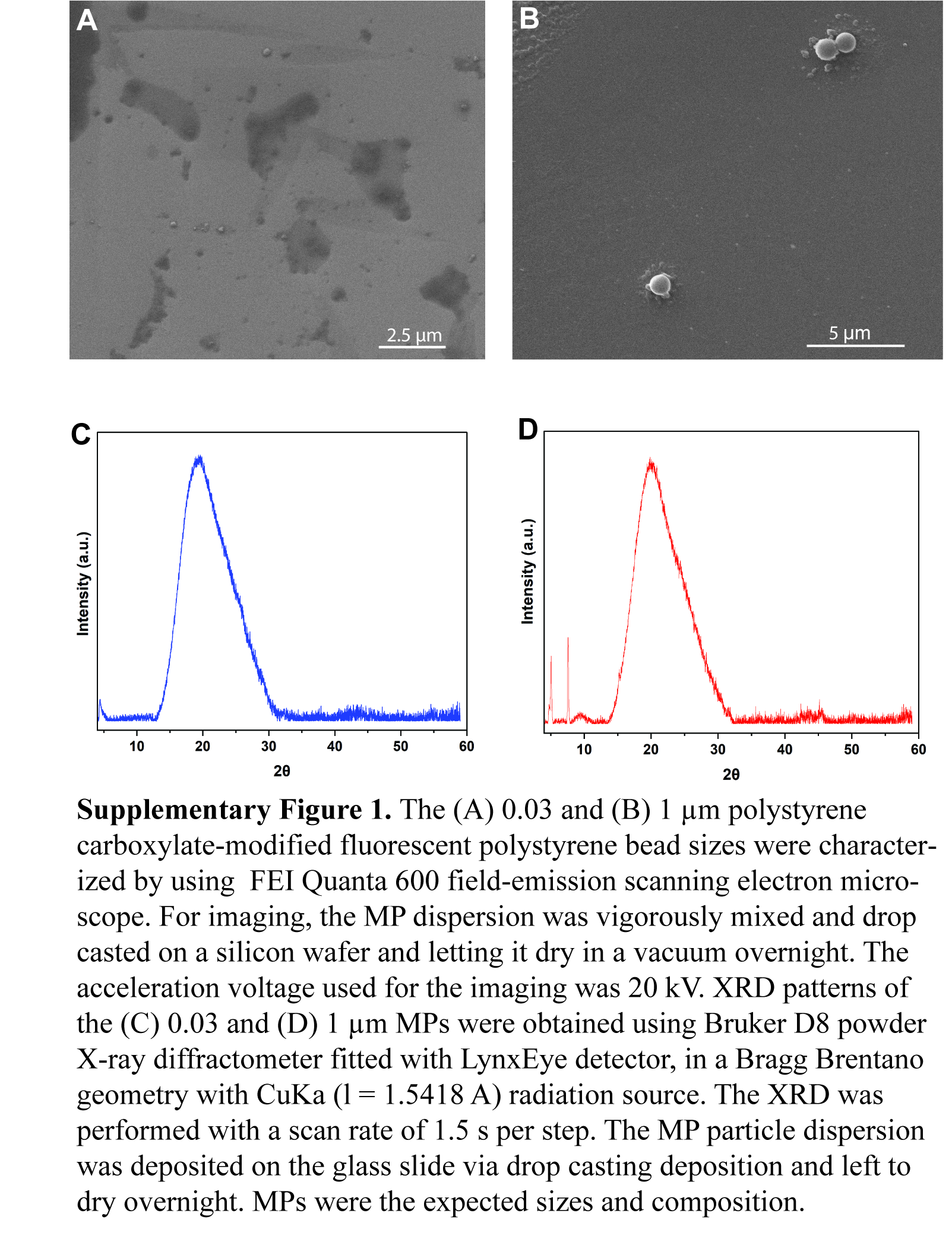
